# Targeting Adaptation to Cancer Treatment by Drug Combinations

**DOI:** 10.1101/2021.04.14.439861

**Authors:** Heping Wang, Augustin Luna, Gonghong Yan, Xubin Li, Ozgun Babur, Gordon B. Mills, Chris Sander, Anil Korkut

## Abstract

Adaptation of tumors to therapeutic interventions contributes to dismal long-term patient outcomes. Adaptation to therapy involves co-action of functionally related proteins that together activate cell survival programs and compensate for the therapeutic impact. Oncogenic dependencies to such adaptive events, however, can generate new therapeutic vulnerabilities that can be targeted with drug combinations. The precision medicine approaches in which targeted drugs are matched to pre-existing genomic aberrations fail to address the adaptive responses and resulting vulnerabilities. Here, we provide the mathematical formulation, implementation and validation of the TargetScore method. The TargetScore identifies collective adaptive responses to targeted interventions as concurrent changes of phospho-proteins that are connected within a signaling network. Based on the adaptive responses, the method predicts drug-induced vulnerabilities. Using TargetScore, we inferred the adaptive responses with short-term (i.e., days) stress and long-term (i.e., months) acquired resistance to inhibitors of anti-apoptotic mediators, MCL1 and BCL2. With experiments guided by the predictions, we identified synergistic interactions between inhibitors of PARP, SHP2, and MCL1 in breast cancer cells. TargetScore is readily applicable to existing precision oncology efforts by matching targeted drug combinations to emerging molecular signatures under therapeutic stress.

## INTRODUCTION

Targeted therapies have led to significant improvements in patient survival in diverse cancer types (1-3). Resistance to targeted therapies, however, is virtually inevitable and can manifest as a lack of initial response to therapy (intrinsic resistance) or disease progression after the temporary response (acquired resistance) (4). Drug combinations can overcome or prevent the resistance by blocking therapy escape routes and can lead to overall survival gains (5-6). However, the discovery of effective combination therapies is a daunting task due to the complexity of the molecular landscapes associated with drug response and resistance.

A recurrent mechanism of resistance is adaptation to the therapeutic stress through activation of oncogenic programs that compensate for the effects of the therapy (5,7). Adaptive responses to therapy can be manifested within days as a consequence of the plasticity of the processes that mediate cell survival (8-9). The long-term adaptive changes, which lead to acquired resistance in time-scales of months after effective therapy, may be a consequence of emerging genetic alterations and yet still be manifested as mechanisms similar to short-term events (e.g., phenotypic shift, differentiation and signaling plasticity) (1-3). These adaptive responses, regardless of their time-scales, may produce new oncogenic dependencies and resulting therapeutic vulnerabilities that can be targeted with a second drug (10). Applications guided by this concept have enabled the preclinical development of a large repertoire of targeted drug combinations. The targets of such drug combinations involve tumor growth, survival, DNA repair, immune evasion, epigenetic regulation, apoptotic mediators and other mechanisms (11-22). Clinical trials guided by adaptive responses are also emerging as exemplified by co-targeting PARP with signaling and DNA repair pathways (23-24), and in general, clinical trials guided by biomarkers increase the success rate of trials at all stages (25). However, the adaptive response mechanisms and resulting therapeutic opportunities are not fully explored owing to the number of combinations possible in diverse cancer contexts. Moreover, it is highly challenging to prioritize effective combination therapy targets among the broad set of oncogenic processes that can be affected by a first-line therapy.

Analytical methods that extract oncogenic dependencies from high throughput data may accelerate the implementation of combination therapies. Methods, such as network modeling, that capture collective behaviors of functionally associated molecules may provide stronger predictions compared to methods that focus on isolated processes (26). The use of phospho-proteomic data improves the prediction of responses to targeted drugs since targeted agents usually act by altering the post-translational modifications of oncogenic proteins (27-28). Additionally, computational methods that can generate testable predictions based on readily available (or easily generated) data can be more widely disseminated and have a broad impact on precision oncology efforts (29). A recently reported method, DrugCell involves the implementation of neural networks represented with complex hierarchical structure of a cell and training with responses to hundreds of drugs in approximately one thousand cell lines (30). In the absence of protein activity data, another approach, OncoTreat, relies on solving the highly challenging task of predicting protein activity from mRNA expression data to prioritize drugs based on their ability to suppress the activity of key oncogenic proteins (31). Logical models (e.g., CellNOpt) can be trained on a combination of prior signaling information and comprehensive proteomic drug response data to predict unseen drug responses using either Boolean or differential equation formalism (32). And similarly, our Perturbation Biology method predicts responses to unseen perturbations using interpretable network models inferred with statistical physics algorithms that require large-scale, comprehensive drug response data (33-34). Despite the significant contributions to automating precision therapy selection, a majority of the previous approaches (i) focused on transcriptional or mutational changes, (ii) required comprehensive datasets as modeling constraints that are costly to generate, and (iii) cannot be immediately scaled to a large number of biological samples or molecular entities due to requirement of highly comprehensive datasets.

We have developed a statistical network modeling method, TargetScore to predict effective combination therapies. Based on phospho-proteomic responses to only a few perturbations, TargetScore quantifies the collective adaptive resistance responses within signaling networks. The drug combinations that are predicted to suppress the adaptive responses are tested experimentally and validated combinations are nominated for clinical trials. Using the TargetScore, we analyzed the responses in breast and ovarian cancer cells to inhibitors of anti-apoptotic proteins, MCL1 and BCL2. We quantified adaptive responses under short-term therapeutic stress and with long-term therapeutic exposure leading to acquired resistance. We showed that the emergence of acquired resistance to MCL1 and BCL2 inhibitors is accompanied by increased levels of cell growth (HER2/GAB2/SHP2/MAPK) and DNA repair markers (poly (ADP-ribose) (PAR)/PARP). Drug combinations involving pairs of MCL1, PARP, and SHP2 inhibitors were synergistic and effective in varying contexts of BCL2/MCL1 inhibitor resistance. As demonstrated in models of resistance to apoptotic targeting, TargetScore enables the development of effective combination therapies that target adaptive responses. The combination therapies from TargetScore may provide durable responses in patients that carry the adaptive response signatures.

## RESULTS

### TargetScore algorithm quantifies collective responses to targeted perturbations

The TargetScore algorithm quantifies adaptive responses to perturbations and nominates rational combination therapies to suppress the adaptation mechanisms (Figure 1). The algorithm is built on the observations that (i) cancer cells adapt to the therapeutic impact of drugs through activation of compensatory survival and proliferation programs (Figure 1A) (5); (ii) the compensatory programs depend on the co-activation of functionally related molecules (e.g., members of a signaling pathway) that together drive the adaptation (35,26,12). A corollary to the the observations is that concurrent activation of functionally linked proteins in response to therapy is a more likely predictor of adaptation compared to the responses of individual molecules (Figure 1B). The TargetScore algorithm searches for such “collective adaptive responses” to therapy, co-activation of functionally linked oncogenic proteins (or de-activation of tumor suppressors) that lead to drug resistance (Figure 1C). The TargetScore algorithm involves the calculation of a reference network that captures the functional links between signaling molecules followed by quantification of sample-specific adaptive responses carried by proteins that are linked in the reference model (Figure 1D) (See methods section for mathematical formulation and details).

**Figure 1.**
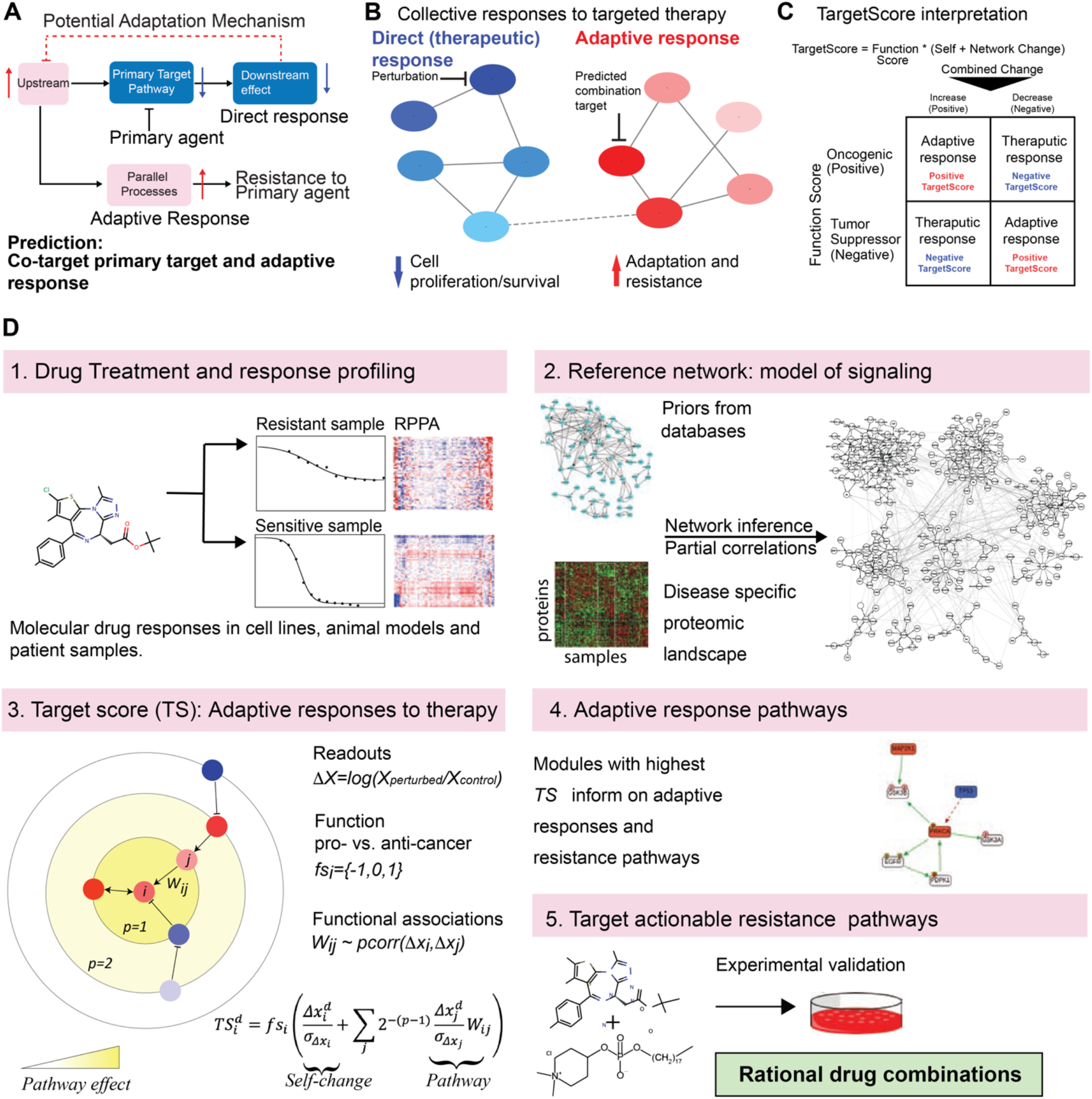
The TargetScore Algorithm. **(A)** Tumors adapt to stress engendered by targeted therapies through activation of compensatory proliferation or survival programs leading to the emergence of adaptive resistance. Adaptive responses can, in turn, be targeted with combination therapies. **(B)** The responses to targeted therapies are, in most cases, collective meaning functionally associated molecules work together (in parallel or sequential in time) to execute a phenotypic shift that enables escape from therapeutic effects. **(C)** TargetScore quantifies collective responses to targeted therapies as a sum of the self-changes in each protein plus the changes in functionally linked entities (i.e., pathway neighborhood). The oncogenic vs. tumor suppressor events are annotated to each protein i with a function score (fs_i_: +1 for oncogenic, -1 for tumor suppressor proteins) **(D)** The TargetScore algorithm steps. 1. Profiling of the responses to a perturbation; 2. Inference of a reference network using the prior-glasso algorithm. Proteomic data from a set of samples that represent a specific context (e.g., disease type) and prior signaling information from databases serve as constraints for network inference (see methods). On the reference network, the signaling interactions between nodes (phospho-proteins i and j) are quantified by the edge strength, W_ij_ (W_ij_ > 0: activating interactions that increase phosphorylation or expression, W_ij_ < 0: deactivating interactions that decrease phosphorylation or expression). 3. Quantification of a sample and context-specific adaptation score (i.e., a TargetScore value) on the reference network using molecular drug response data (See equation 10). 4. Identification of network modules (collective changes) that have a significantly high TargetScore in each sample. The network modules involved in adaptive responses are determined by mapping the TargetScore values back on the reference network and extracting the connected sub-networks enriched with high TargetScore values. 5. Selection of actionable targets that participate in adaptive responses in a sample and testing drug combinations in preclinical models.

The reference network model is inferred based on phospho-proteomics data from a population of samples with shared characteristics. The samples with shared characteristics may belong to a particular cancer type (e.g., breast cancer patients) or a cohort defined by a genomic aberration such as KRAS-mutation. The inter-patient co-variations of the phospho-protein levels, which arise due to perturbations act as the constraints for inference of the signaling network. The perturbations can be intrinsic as mutations and epigenetic changes or extrinsic as drug treatments. In model inference, we also benefit from prior signaling information as imported from the signedPC resource within the Pathway Commons signaling database (44). In the network models, the nodes represent the phospho-proteomic measurements and edges represent the direct associations between the nodes after removal of confounding factors from other variables. The rationale in the model inference is that if the levels of two phospho-proteins are directly associated with each other across diverse perturbations, the two proteins likely function together and are connected to each within the network. For inference, we selected the graphical LASSO (glasso) algorithm based on our comprehensive benchmarking of network inference methods for proteomics-based modeling of signaling interactions (36). The glasso is a sparse penalized maximum likelihood estimator for the inverse of a covariance matrix and used to infer partial correlation-based networks (37). To incorporate the prior signaling information, we generated a modified glasso algorithm, termed prior-glasso that introduces priors as a probabilistic bias similar to the complexity term in the log-likelihood equation (Equation 4 in Methods). The prior interactions that conform to the data are favored, priors that do not conform with data are eliminated and new interactions are inferred solely from data. The resulting network model provides a map of functionally linked proteins on which sample-specific adaptive responses can be quantified.

Based on the interactions within the reference network and sample-specific phospho-proteomic drug response data, the adaptation to targeted agents is quantified by the TargetScore (Equation 10). The TargetScore values are calculated as the network interaction weighted sum of the “self-change” of each protein and the change in its pathway/network neighborhood on the reference network in response to targeted perturbations. High TargetScore values correspond to adaptive responses (e.g., increased RTK activity by MEK inhibitor via a feedback loop, (38) and low values correspond to the therapeutic impact of the drug (e.g., lowered p-ERK in response to MEK inhibition) (Figure 1C). The connected nodes with high TargetScores form the drug-activated network modules of signaling and are predictors of collective adaptive resistance mechanisms. The predictions from the algorithm are effective drug combinations that target and suppress adaptive responses. When short term adaptive-responses are targeted, the combination may involve the first agent that induces and the second agent that suppresses the adaptive response. In the case of acquired resistance, a potentially more effective strategy is to target the multiple adaptive events as cells are likely hard-wired in the long-term through genetic or epigenetic changes to gain resistance to the first agent. The comparison of TargetScore values over different cancer samples (e.g., sensitive vs. resistant) reveals how drugs differentially rewire pathway activities to quantify adaptive events. The calculations with samples on varying treatment time scales (e.g., days to months) inform on the evolution of drug resistance. In summary, with a selection of versatile agents, doses, timeframes and samples; the algorithm explores adaptation to therapy across diverse dimensions (e.g., drug targets, timescales, degree of resistance). The predicted drug combinations are tested experimentally and nominated for further preclinical development and clinical trials.

### Emergence of resistance to apoptotic targeting in MCL1-amplified breast cancer cells

We developed experimental models of resistance to MCL1 and BCL2 inhibitors using the HCC1954 breast cancer cell line (Figure 2A). The MCL1 amplified HCC1954 cell line is highly sensitive to MCL1 inhibitors (MCL1i; Figure 2B) and partially sensitive to BCL2 inhibitors (BCL2i; Figure 2C). We generated the cell lines, named HCC1954-MR and HCC1954-BR, with acquired resistance to MCL1i (S63845) and BCL2i (ABT-199), respectively, through the long-term treatment of each agent in HCC1954 cells over the course of 4 months with increasing doses (0 to 4μM) (Figure 2B). As a model of intrinsic resistance to MCL1 inhibition, we used the SKOV3 ovarian cancer line, which is diploid for MCL1 and relatively refractory to MCL1 inhibition. The selection of the SKOV3 cell line is further justified as it is a model of high-grade serous ovarian cancer, a subtype with high molecular and pharmacological similarity to basal-like breast cancers, which is the origin of HCC1954 (39).

**Figure 2.**
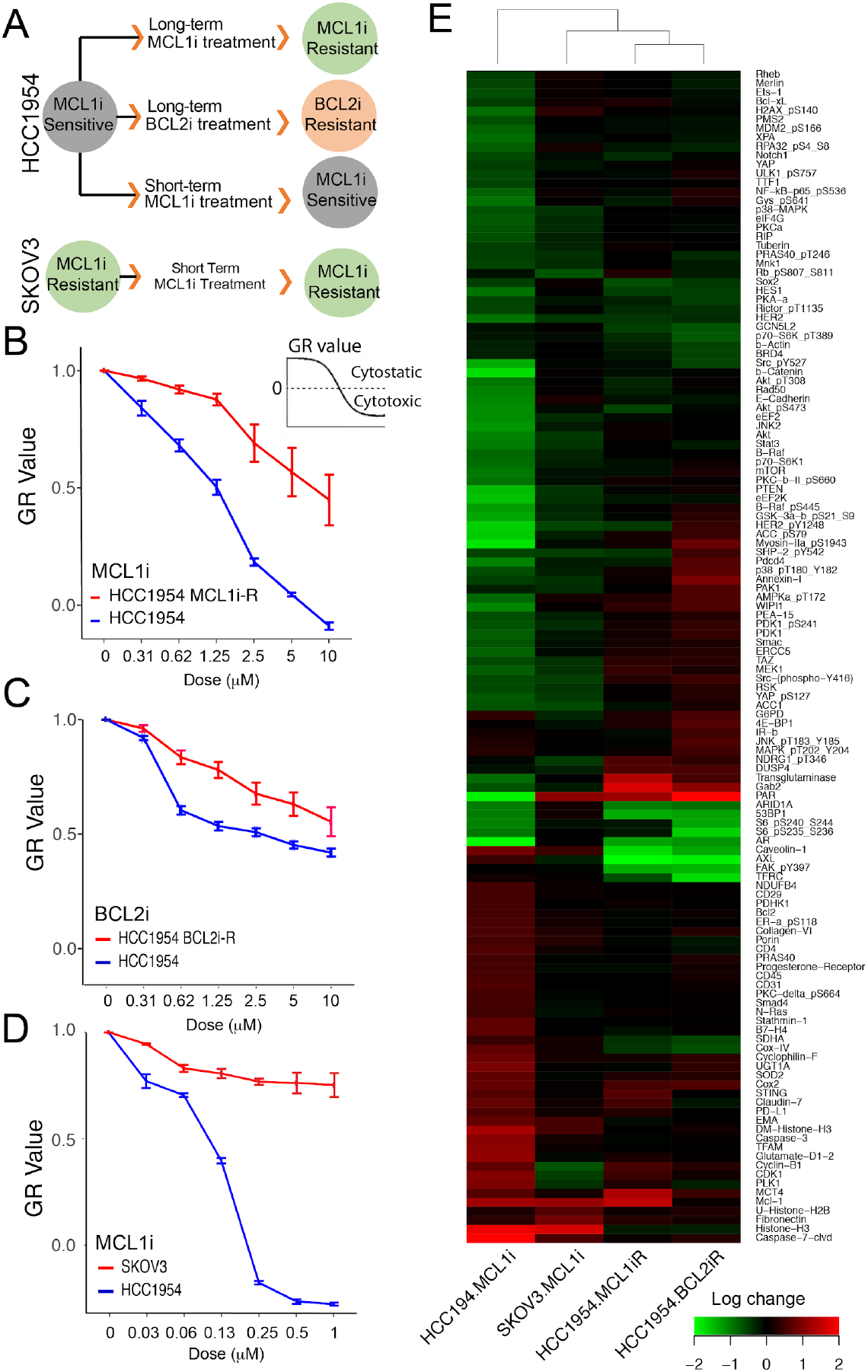
Response and resistance to apoptotic agents in cancer cells. (A) Generation of model systems with acquired and intrinsic resistance to MCL1i and BCL2i. The HCC1954 breast cancer cells (MCL1 amplified) were treated with MCL1i or BCL2i at longitudinally increasing doses until the emergence of resistance (∼4 months). SKOV3 cells (MCL1 wild type) represent a model with intrinsic MCL1i resistance. (B) Growth rate (GR) changes in response to MCL1i in cells with acquired resistance to MCL1 targeting. In cells with acquired MCL1i resistance, (HCC1954-MR), response is measured 3 days after multiple doses of acute MCL1i treatment. GR 0 to 1: partial growth inhibition (cytostatic); GR 0 to -1: cell death (cytotoxic). (C) Growth rate changes in response to BCL2i in cells with acquired resistance to BCL2i. (D) Responses to MCL1i in cells with SKOV3 vs. HCC1954 cells. (E) Phospho-proteomic profiling of responses to short and long-term MCL2i or BCL2i treatment in cells with varying drug resistance. The phospho-proteins whose expression changes in response to at least one perturbation are included (max (abs(log2(x_ik_/x_io_)) > 0.5, for protein i across all perturbations k; x_ik_: readout in perturbed state; x_io_: readout in unperturbed state).

To monitor drug response, we used the growth rate (GR) inhibition metric (40). GR mitigates the confounding factors from varying duplication times across different cell lines. This is particularly critical for the analysis of acquired resistance as the emergence of resistance generally alters cellular doubling times (41-42). In HCC1954 cells, the doubling time was increased from a baseline of 44 hours to 73 and 79 hours in HCC1954-MR and HCC1954-BR, respectively (Figure S2A). Using Reverse Phase Protein Array (RPPA) based measurements, we profiled the changes in >300 oncogenic protein levels and phosphorylation states in HCC1954, HCC1954-MR, HCC1954-BR and SKOV3 in response to apoptotic perturbations (Figure 2C, S2B). The parental HCC1954 and SKOV3 are molecularly profiled with short-term drug treatment (48 hours), while HCC1954-MR and HCC1954-BR are profiled after long-term drug perturbations (4 months). The proteomic data (Figure 2E) covered diverse oncogenic processes including PI3K, RAS-MAPK, Src/FAK, TGF signaling axes, DNA repair, cell cycle, apoptosis, immuno-oncology, metabolism enzyme levels, and histone modifications. This resource on BCL2/MCL1 inhibitor responses and resistance modalities served as the input data for the TargetScore analysis.

### TargetScore identifies adaptive responses to apoptotic targeting

We applied the TargetScore algorithm to investigate the collective adaptive responses to apoptotic targeting in each cell line with varying resistance to MCL1 and BCL2 inhibitors. First, we inferred the disease-type specific reference networks for breast and ovarian cancers. Next, we calculated the sample-specific and network-level responses to inhibitors of anti-apoptotic proteins, MCL1 and BCL2 for cells with varying drug resistance.

The reference models are inferred using 1) RPPA data from breast and ovarian cancer TCGA cohorts and 2) prior knowledge interaction information (Figure 3, Figure S3) (39, 43-44). The combination of these two data sources enables consistency with both context-specific biological characteristics (e.g., resulting from the cancer types studied) and previously reported signaling knowledge. The breast and ovarian cancer datasets cover phospho-proteomic data from 901 and 408 patients respectively. Both datasets covered 224 phospho-protein species. The network models are inferred with the prior-glasso method (see Methods for mathematical formulation). To enable comprehensive adaptive response quantification in subsequent steps, the interactions involving the proteins that are not covered by the patient RPPA but measured within the drug response data are also included solely based on prior information. The model complexity and prior weights in the prior-glasso inference framework (equation 4) are optimized to maximize the glasso likelihood function with a BIC calculation (Equation 5) and set as ρ=0.035, κ=0.025 for the breast cancer model (Figure 3B). The ovarian cancer model is inferred using the identical approach (Figure S3B). We then tested the model robustness through bootstrapping partial datasets (1000 bootstraps, see methods) with varying data coverage (50% to 90%) and comparison of models based-on partial data to models inferred from the complete data (Figure 3C). Even in the lowest coverage level (50%), the models from partial data had a median of 80% overlap with models from the complete data suggesting a highly robust inference. Next, the model accuracy was tested with 5-fold cross-validation with 500 bootstraps (equation 9), demonstrating models inferred with actual data reach significantly lower errors than label-randomized data (Figure 3D). For the breast and ovarian cancer signaling models, the resulting reference networks carry 954 and 1017 edges (W_ij_ > 0.1) respectively, connecting 304 nodes, with a varying degree of 1 to 40 edges/node (Figure 3E, S3C). The disease-type specific reference networks (Figure 3F, S3) served as the reference models to calculate the sample-specific adaptive responses.

**Figure 3.**
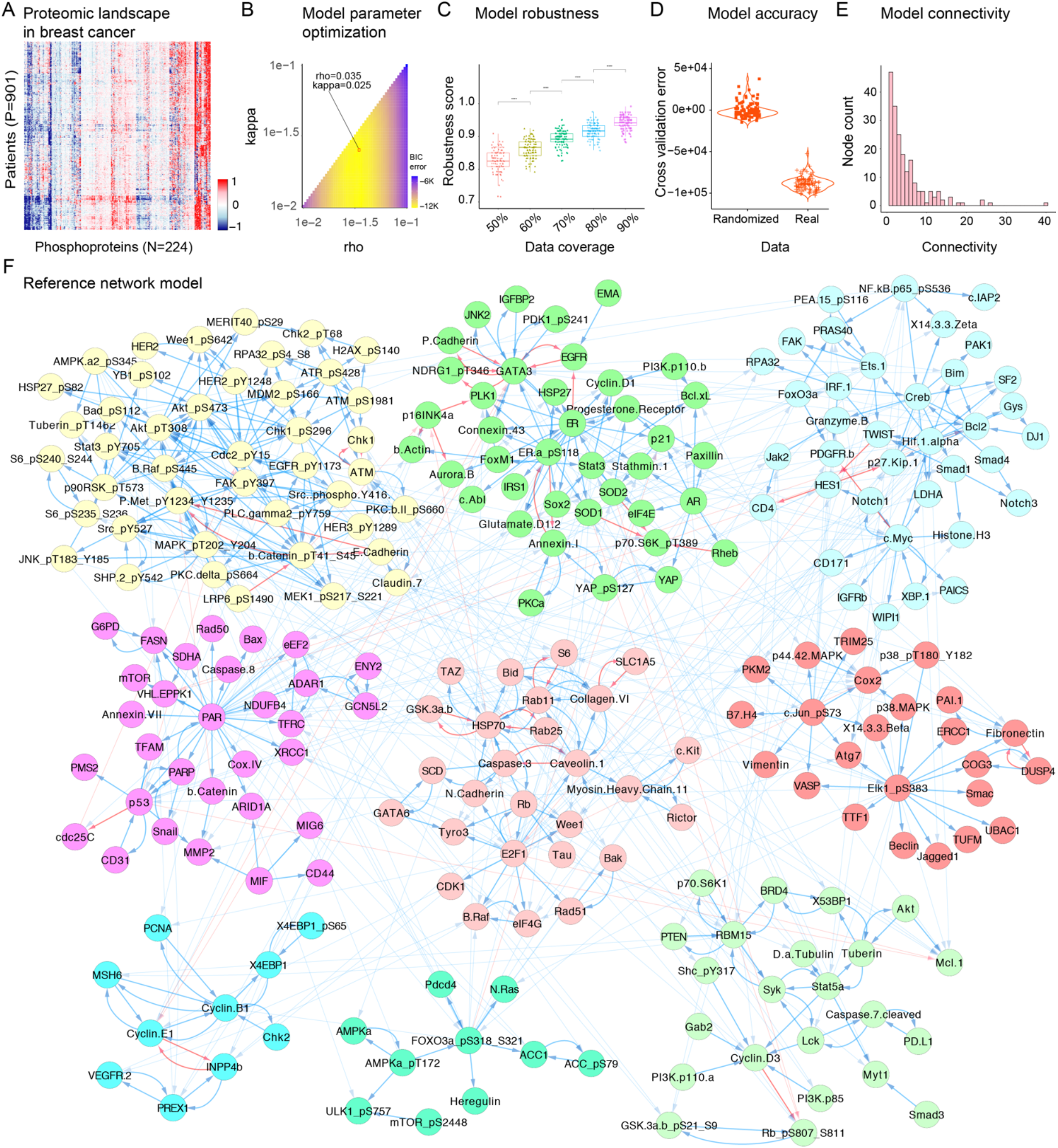
Reference network captures disease type-specific phospho- and total proteomic associations on which collective adaptive responses can be quantified. (A) The breast cancer phospho-proteomics data for cancer-relevant proteins from 901 patients. (B) Optimization of prior-glasso algorithm parameters is based on BIC error across all *ρ* (model complexity penalty) and *κ* (prior information prize). (C) The robustness of the reference network is quantified by the concordance between network models that were calculated with complete vs. subsampled partial data with varying coverage (50% to 90%). (D) Network model validation. The comparison between errors of models that were inferred using actual RPPA versus label-randomized data. (E) The distribution of connectivity in the network models. (F) The reference network model of signaling interactions for breast cancer is inferred using the prior-glasso algorithm. Blue edges represent activating and red edges represent deactivating interactions. The network modules are calculated using the GLay community clustering method based on the Newman-Girvan algorithm and each module is represented by a distinct color (45). Interactions with W_ij_ > 0.1 are shown.

We calculated the sample-specific TargetScores with reference models and phospho-proteomic drug response data from HCC1954, HCC1954-BR, HCC1954-MR and SKOV3 lines treated with BCL2 and/or MCL1 inhibitors. The calculations for HCC1954 and SKOV3 cells utilized the breast and ovarian cancer reference networks, respectively. A TargetScore value is calculated for each measured molecular entity (here either a protein or phospho-protein) for all perturbation conditions in all cell lines with varying sensitivities to apoptotic targeting (Equation 10). An FDR-adjusted p-value is calculated for each TargetScore value based on a null distribution of scores from randomized drug response data (see Methods). The resulting TargetScores have provided a comprehensive resource to analyze the adaptive responses to apoptotic targeting.

### TargetScore algorithm predicts actionable adaptive responses in DNA repair and growth pathways

We analyzed the TargetScore values for all proteins to identify potential drivers of adaptation to apoptotic targeting with a focus on MCL1 inhibition as MCL1 amplifications are common in breast and ovarian cancers (46). We analyzed the adaptation profiles in the cells with high sensitivity, intrinsic resistance and acquired resistance to apoptotic targeting. In order to nominate potentially effective combination therapies to overcome resistance, we evaluated the TargetScore ranking, statistical significance and differential characteristics across samples with varying resistance to MCL1/BCL2 inhibition Based on ranking and statistical assessment (see Methods) of the TargetScore values, we first focused on the proteins that are particularly involved in acquired resistance to MCL1 and BCL2 targeting (Figures 4B-F). In models of acquired resistance to BCL2 or MCL1 targeting (Figures 4E-F), we observed a likely DNA damage response (DDR) marked by concurrent TP53BP1 depletion (increased Target Score, fs_TP53BP1_=-1) and increased PARylation (labeled PAR) a marker of PARP activity (Figure 4E-F) (47). TP53BP1 depletion can indicate homologous recombination (HR) mediated double break (DSB) repair, while PARylation may be indicative of single-stranded break (SSB) repair driven by PARP enzyme activity (48,49). In parallel, GAB2 adapter protein, which links receptor tyrosine kinases to downstream SHP2/MAPK growth signaling (50,51), had significantly high TargetScore values in the HCC1954-MR and -BR acquired resistance models (Figure 4E-F).

**Figure 4.**
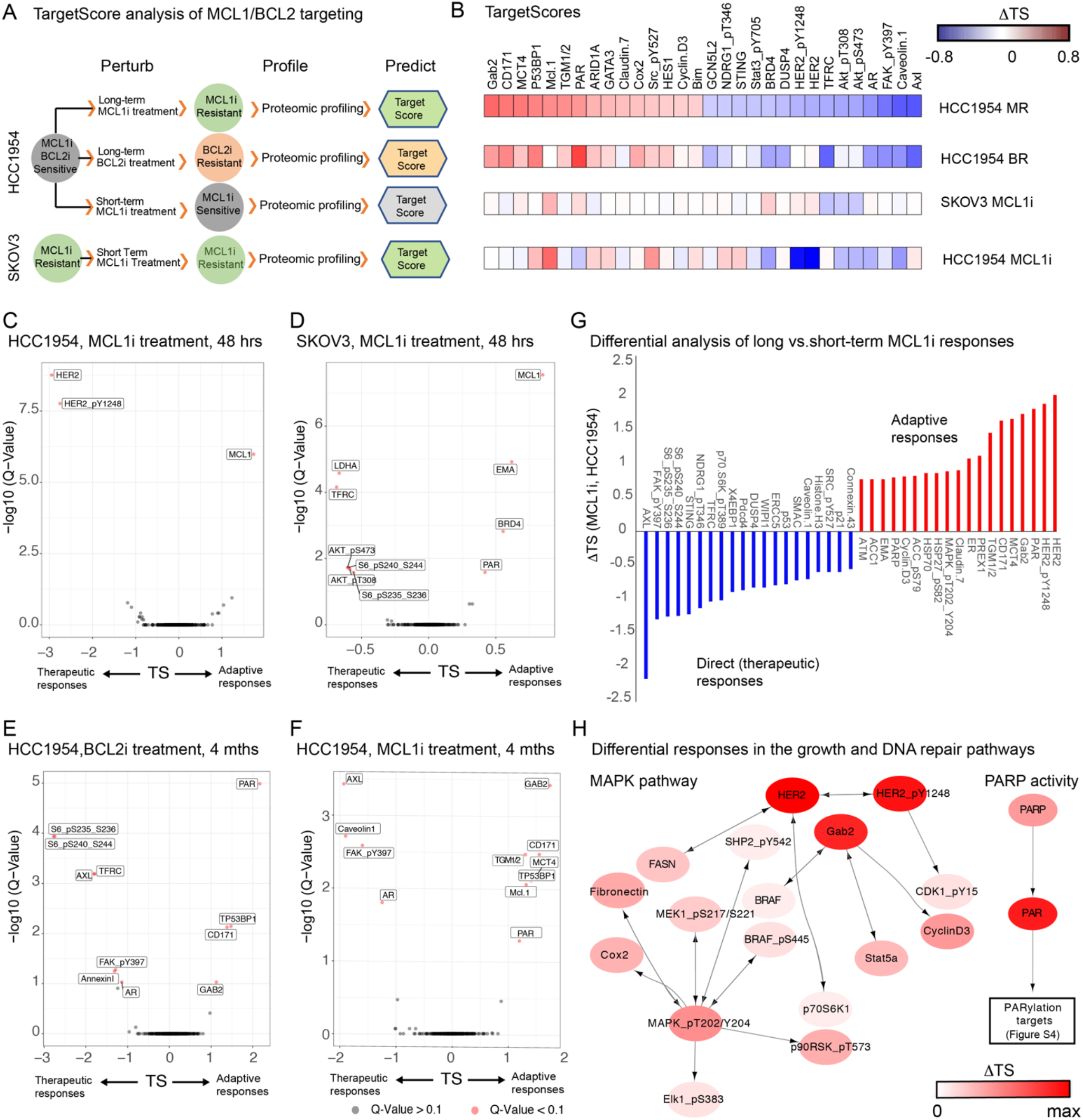
TargetScore algorithm nominates maps of adaptive responses and targets to mitigate drug resistance. (A) The proteomic profiles from drug-resistant models serve as constraints in TargetScore calculations. (B) The proteins are ranked based on the highest and lowest TargetScore values in cells with acquired resistance to MCL1i. (C-F) The statistical assessment of TargetScores for cells with varying MCL1i/BCL2i resistance. The P-values are based on a null distribution generated from randomized data and corrected with FDR-adjustment (Benjamini-Hochberg method). (G) The differential analysis of TargetScores upon short- and long-term exposure to MCL1i in HCC1954. The highest positive (red bars) and negative (blue bars) changes are shown based on the difference of TargetScores from resistant vs. sensitive cells. (H) The network representation of the differential responses in short vs. long-term MCL1i exposure in MAPK and PARP networks (Figure S4 for downstream PARylation targets).

Next, we addressed the distinct characteristics of long-term responses to MCL1 targeting. We performed a differential analysis of TargetScores from HCC1954 cells with short-term (48 hours) vs. long-term (HCC1954-MR, 4 months in increasing doses) exposure to the inhibitor (Figure 4G). This analysis confirmed the involvement of growth and DNA repair pathways (Figure 4H). In addition to increases in GAB2, acquired resistance is marked with the restoration of other RTK/MAPK pathway mediators including the actionable HER2/p-HER2, SHP2, RAF/MEK/ERK as indicated by differential analysis of TargetScores from MCL1i sensitive and acquired resistance models (Figure 4H). Similarly, levels of both PARP and the PARylation markers that are indicative of PARP activity are selectively high in samples with acquired resistance to MCL1 inhibition suggesting a potential dependence on double-stranded DNA-repair (Figure 4H).

In addition to the aforementioned DNA repair and growth pathway markers, we observed significant increases in the cell adhesion molecule CD171 (L1CAM) as well as decreases in androgen receptor (AR), receptor tyrosine kinase AXL, and p-FAK in both BCL2i (Figure 4E) and MCL1i (Figure 4F) resistant models. With potential relevance to drug resistance, CD171/L1CAM has already been implicated in resistance to apoptosis and poor outcome in various cancers as well as epithelial to mesenchymal transition (EMT) (52). However, we did not track CD171 and potential EMT responses due to the challenges associated with the therapeutic targeting of EMT. In the de novo resistance model, SKOV3 cells, we observed concurrent high TargetScores for BRD4 and PARylation (Figure 4D). We have previously demonstrated co-targeting BRD4 and PARP is highly synergistic in diverse in vitro and vivo models including the SKOV3 cell line (16). The MCL1 TargetScore value was also high in all MCL1 inhibitor-treated samples regardless of their level of sensitivity to MCL1 inhibition (HCC1954, SKOV3, HCC1954-MR) suggesting that an increase in MCL1 is not unique to drug-resistant context and does not cause resistance to MCL1 inhibition (Figure 4C-E). Consistent with previous studies, the increase in MCL1 protein and associated high TargetScore upon MCL1 inhibition is likely due to stabilization of the inhibited protein by drug binding and has no anti-apoptotic consequence (53).

Based on the consistently high TargetScore values associated with clinically relevant mediators of DNA repair and cell growth pathways, we turned our attention to the therapeutic targeting of these two key oncogenic processes.

### Co-targeting PARP and SHP2 is synergistic in cells with resistance to apoptotic targeting

Guided by the TargetScore predictions, we experimentally measured the impact of co-targeting DNA repair, growth pathways and the anti-apoptotic mediators particularly in the drug-resistant cells. For this purpose, we treated the HCC1954, HCC1954-MR and HCC1954-BR cells with paired drug combinations (Figure 5). When possible, the proteins were directly targeted with specific inhibitors as in the case of PARP, MCL1 and BCL2. On the other hand, the GAB2 molecule is not directly actionable with existing drugs. Therefore, we chose the GAB2 interaction partner, SHP2 phosphatase that links GAB2 to MAPK signaling (54). SHP2 inhibitors have been shown to disrupt signaling downstream of GAB2 (e.g., MAPK signaling) (54,55,56) by decreasing the interaction of SHP2 and GAB2 (57). We quantified responses using the growth rate (GR) metric (Figure 5A-C). The drug synergy is quantified using the combination index (CI) based on the IC50 doses of each agent (CI < 0.8: synergy; 0.8 < CI < 1.2: additive; CI > 1.2: antagonism) (Figure 5D).

**Figure 5.**
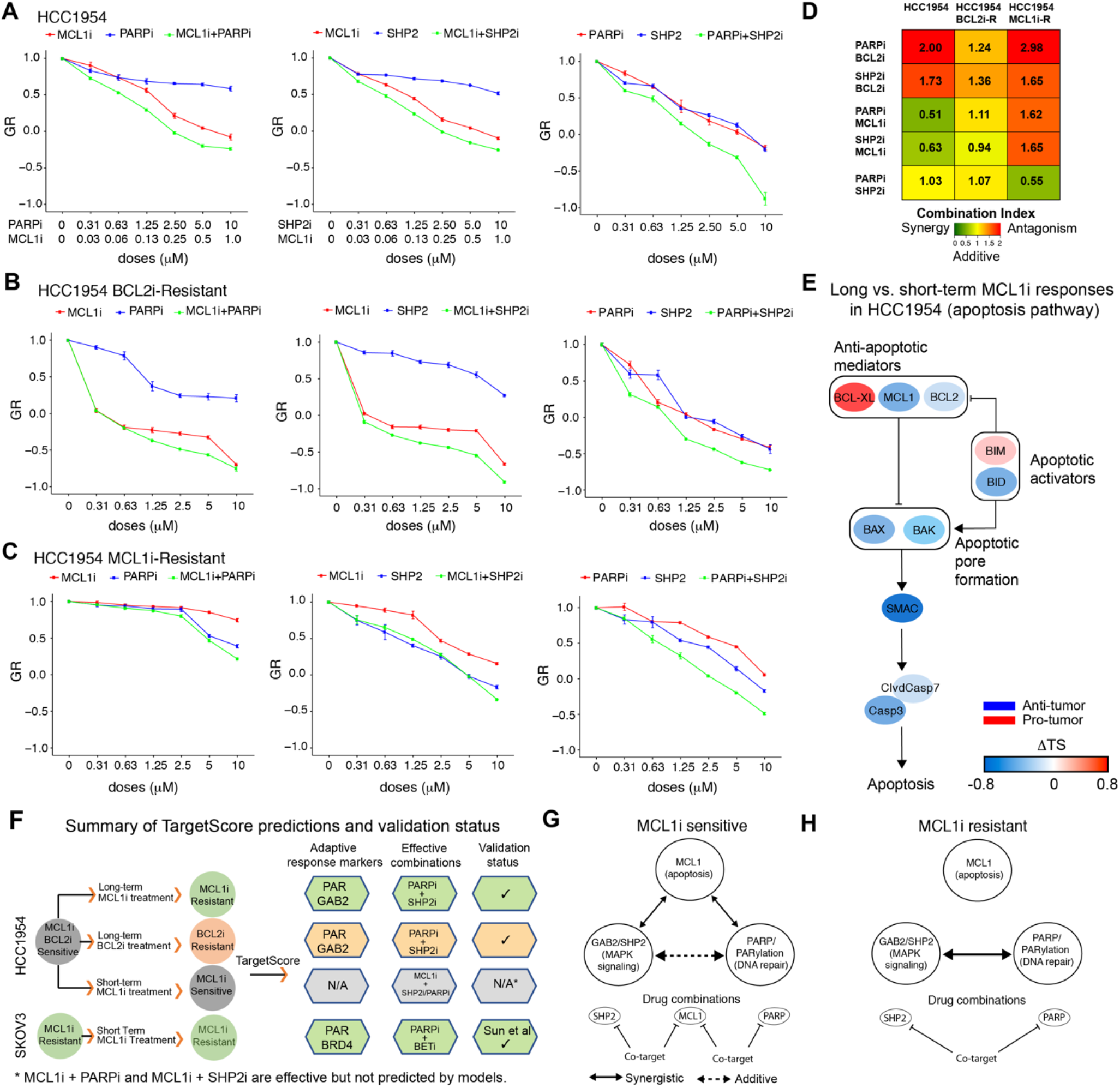
Rational combination therapies from TargetScore analysis. Growth rate changes in response to combinations of PARP, SHP2, and MCL1 inhibitors in (A) HCC1954; (B) HCC1954 resistant to BCL2i; (C) HCC1954 resistant to MCL1i (Figure S5 for cell viability changes and combinations with BCL2i). (D) The combination indexes quantify the synergistic drug interactions at IC50. (E) The differential analysis of TargetScores within the intrinsic apoptotic pathway in cells with acquired resistance (HCC1954-MR) vs. sensitivity (HCC1954) to MCL1i. The proteins that are covered in the RPPAs and TargetScore are shown. (F) The predictions and experimental testing based on the TargetScore. (G-H) A proposed model of changing interdependencies with the emergence of resistance to MCL1i. In the sensitive cells, the coupling between MCL1, PARP and SHP2 dependencies is reflected in the synergies of co-targeting MCL1+PARP and MCL1+SHP2. In cells with acquired resistance, the synergistic interactions between MCL1 and PARP or SHP2 are lost. The synergy between PARP and SHP2 inhibition, however, has become stronger leading to a potential therapeutic benefit in the resistant cells. Solid lines: interactions that lead to synergistic drug responses, dashed lines: interactions with additive responses.

First, we focused on the combined inhibition of the targets that were increased in cells with acquired resistance to MCL1 and BCL2 inhibition. As predicted by the high PARylation and GAB2 TargetScores (Figure 4H), co-targeting PARP and SHP2 in HCC1954-MR (acquired resistance to MCL1 inhibitor) was highly synergistic (CI=0.55, Figure 5D) and cytotoxic effects were reached at relatively low doses (2.5μM) (Figure 5C). In HCC1954-BR (acquired resistance to BCL2 inhibitor), a considerable cytotoxic effect is reached even with the single-agent MCL1 inhibitor treatment (Figure 5B), and no strong improvement with the addition of either PARP or SHP2 inhibitors is observed (Figure 5B,D). In line with the predictions (Figure 5F), the HCC1954-BR cells also had increased sensitivity to PARP and SHP2 co-targeting (cytotoxic at 0.625μM, Figure 5B) with an additive interaction (CI=1.07, Figure 5D).

In HCC1954-MR cells, co-targeting of MCL1 with either SHP2 or PARP did not introduce any benefit (Figure 5C). Both combinations were antagonistic (CI > 1.5, Figure 5D) and cytostatic effects were not observed at relevant doses. With a retrospective differential analysis of the TargetScore values in resistant vs. sensitive parental cells, we observed that the BCL-XL TargetScore was increased as resistance to the MCL1 inhibition emerged (Figure 5E). This observation leads to the prediction that anti-apoptotic dependence is shifted from MCL1 to BCL-XL activity concurrently with loss of responsiveness to MCL1 targeting. Indeed, it was recently reported that small cell lung carcinoma cells with high MCL1 and low BCL-XL are significantly sensitive to MCL1 targeting and patient cohorts with high MCL1/BCL-XL ratios may be candidates for MCL1 inhibitor therapy (58) (PMID: 32152266). Here, we make the reciprocal but equivalent observation that the emergence of MCL1 inhibitor resistance is accompanied by a decreased MCL1 to BCL-XL ratio. Although the shift in the MCL1 to BCL-XL ratio may point to a relevant adaptation mechanism to chronic suppression of MCL1, we did not experimentally test BCL-XL inhibitors due to their severe toxicity on platelet cells in patients and the reduced enthusiasm for clinical translation (59). On the other hand, the parental sensitive cells (HCC1954) displayed a strong response and synergy for the combinations of MCL1 inhibitor with both PARP and SHP2 targeting (CI(MCL1i-PARPi)=0.51, CI(MCL1i-SHP2i)=0.63) (Figure 5D). Similarly, co-targeting PARP and SHP2 led to strong responses with a CI in the additive range and a shift from cytostatic to cytotoxic effects between 1.25 and 2.5μM in HCC1954 (Figure 5C). This was interesting as the modeling in sensitive cells did not immediately capture such a prediction, suggesting there is an inherent co-dependency on PARP and SHP2 even before the emergence of resistance.

Compared to the MCL1 inhibitor, the BCL2 inhibitor had a limited impact on growth rate as well as viability (Figure S5). The relatively limited efficacy of targeting BCL2 is expected as drug naïve HCC1954 cells are copy number-amplified for the MCL1 gene and have higher baseline MCL1 protein expression, resulting in a potential dependence on the anti-apoptotic activity of MCL1 (53, 71).

In summary, our results suggest the MCL1 inhibition is synergistic with SHP2 and PARP targeting in HCC1954 cells (Figure 5G, H) and co-targeting SHP2 and PARP can generate significant therapeutic benefit with increased synergy in MCL1i and BCL2i resistant cells as predicted by the network modeling (Figure 4).

## DISCUSSION

In the conventional precision medicine paradigm, targeted therapies are selected to suppress the effects of pre-existing oncogenic aberrations, in most cases with mono-therapies that are matched to an individual genomic aberration. Although genomically-matched monotherapies have introduced substantial therapeutic benefits in select cancer types (60, 61,62), responses to targeted therapies are not durable and eligible patient cohorts are limited. As of 2019, only 15% of cancer patients are eligible for genomics-informed therapy and only 8% of patients benefit from such therapies with objective, but nevertheless, transient responses (63).

Targeting adaptation to first-line cancer treatments may improve response duration and depth in larger patient populations. Here, we developed the TargetScore algorithm for better identification of adaptive response mechanisms to targeted agents. We have demonstrated its utility in the discovery of combination therapies associated with response and resistance to targeting anti-apoptotic mediators. The algorithm is built on the rationale that adaptive responses under therapeutic stress occur through protein network rewiring, and collective changes in pathway activities are predictors of mitigation strategies. There have been similar approaches on modeling to predict responses to therapy based on mRNA data (64) or protein-based inference of mainly descriptive network models (65). The predictive proteomics-based network modeling approach has a higher chance of success as drug responses are mediated primarily by functional proteomic changes.

TargetScore has flexible data and computing requirements allowing effective scaling of analyses. The algorithm is built on functions that can support the analysis of both small and large numbers of samples. Therefore, its rapid applications to diverse preclinical or clinical problems is not prohibited. For example, in comparison to our previously published network modeling methods that enable “de novo” drug combination predictions based on rich perturbation response data (33, 34), the minimal molecular data requirement for TargetScore is reduced from hundreds of conditions to a single sample treated with a single agent. Although the TargetScore results are simple to obtain and interpret; here we show that the resulting predictions are sufficient to nominate effective drug combinations. The comparison of TargetScore values over different cancer samples (e.g., sensitive vs. resistant), across different drug doses or time points reveals how drugs rewire pathway activities in time, dose and sample space. The method is amenable to the comparison of pre- and post-therapy samples to quantify both short- and long-term adaptive events; the molecular profiling data needed as input can be obtained from various biological sources: cell lines, xenograft models and patients in clinical trials. The predictions from the method are drug combinations involving the agent that induces an adaptive response and a second agent that interdicts the adaptive response. Future studies involving TargetScore will explore the strength of the methodology utilizing other forms of molecular profiling data (e.g., RNA-seq data) and will seek to couple the TargetScore method with separate methods directed at understanding drug combination toxicities, which also limit the translation of drug combinations to clinical practice (6).

Our analysis suggests that the activity of the MAPK pathway and DNA repair mechanisms were increased in cells with acquired resistance to BCL2/MCL1 inhibitors. In subsequent perturbation experiments guided by the calculations, we observed synergistic and strong responses to combinations of PARP and SHP2 inhibitors in the acquired resistance contexts. In the cells with acquired resistance to BCL2/MCL1 inhibitors, the combinations involving the “first-line therapy” agents (i.e., the MCL1 or BCL2 inhibitor) paired with inhibitors of individual adaptive response markers (PARP or SHP2) were not as effective as a combination composed of PARP and SHP2 inhibitors. In the parental HCC1954 cell line with higher sensitivity to MCL1 inhibition, however, MCL1 co-targeting with either PARP or SHP2 was strong and synergistic (Figure 5D). In this drug sensitive parental line, both PARP and SHP2 targeting was effective yet the drug-drug interactions were more modest (i.e., additive) in comparison to the synergistic effects in the MCL1i resistant cells (Figure 5D). These results suggest, that once resistance is acquired, it is challenging to overcome the acquired resistance to a particular agent (e.g., MCL1 inhibitor) through combinations involving this initial particular agent and an adaptive response marker (e.g., MCL1+PARP or MCL1+SHP2 inhibitors). Nevertheless, a therapeutic benefit from such combinations (e.g., MCL1+PARP or MCL1+SHP2) may be achieved prior to the emergence of resistance. We believe that further study of the observations here is warranted and could be conducted by orthogonal means to assess the clinical utility of these observations. As the BCL2 and MCL1 inhibitors are in clinical trials for the treatment of diverse cancers (66, 67), we expect this study to help guide the selection of clinically relevant combination therapies to target adaptation to apoptotic therapies.

In conclusion, systems biology and bioinformatics analyses of adaptive responses along the therapy timeline may guide the selection of effective combination therapies. Utilization of our analytical approach combined with a longitudinal collection of patient molecular profiling data may lead to a novel precision medicine paradigm based on “matching combination therapies to emerging molecular signatures under therapy stress”. Subsequently, with a broader repertoire of precision combination therapies that are matched to emerging signatures, a larger patient population may benefit from deeper and more durable responses.

## Acknowledgment

This work is supported with grants from MDACC Support Grant P30 CA016672 (the Bioinformatics Shared Resource) (AK), OCRF Collaborative Research Award (AK, GBM), U.S. NCI grants U24CA210950 (GBM), P50CA217685 (GBM), U01CA217842 (GBM), ICI Fund (AK), CPRIT High-Impact/High-Risk Award (RP170640) (AK, GBM), a kind gift from the Miriam and Sheldon Adelson Medical Research Foundation (GBM), SAC110052 from the Susan G. Komen Foundation (GBM), BCRF-18-110 (GBM) and National Institute of General Medical Sciences (P41GM103504) (CS).

## Data sharing statement

The software for the TargetScore algorithm is implemented with an R-based package and the complete source code is available at https://github.com/korkutlab/targetscore. An interactive website for running TargetScore with custom and available datasets is at https://cannin.shinyapps.io/targetscore/ (The tool will be transferred to UT MDACC servers prior to publication). All datasets and perturbations used in this study are shared on the website and ready for TargetScore execution on the website. For rapid testing of the method and software, we also provide a partial dataset covering readouts of 100 phospho-proteins in HCC1954 cells treated with MCL1 inhibitor for 48h in addition to all the complete datasets.

## METHODS

### TargetScore algorithm

#### Data-driven Reference Network Model

The reference network on which the adaptive responses are quantified, captures the signaling interactions between phospho-proteins. In the network models, the nodes represent the levels of phospho-proteins. The edges represent the direct associations between the node pairs while removing out the effects from confounding variables. The reference network is inferred using a data-driven approach with disease or subtype-specific phospho-proteomics data. Based on the data, the models are generated with a statistical inference algorithm, termed prior-glasso and prior information as described below, and briefly in the results section. The interactions within the resulting data-driven model are relevant across the population of samples with shared characteristics.

##### Data sources for network inference

The model inference benefits from phospho-proteomic data covering hundreds of proteomic species across large sample cohorts. The data covers activity markers of key oncogenic signaling molecules as phosphorylation states or total protein levels. The samples are under diverse perturbations (e.g., extrinsic perturbations such as drug treatment or intrinsic perturbations such as mutations/epigenetic states) that create rich information for identifying the molecular associations. Yet, the samples should carry shared characteristics (e.g., disease type, stage, existence of a key mutation) to enable modeling relevant to a specific biological context. The set of phospho-proteins is selected to interrogate major drivers of oncogenic processes and therapeutically actionable targets, leading to biologically relevant models (Table S1). The variations and associations of phospho-protein levels that emerge under different perturbations serve as the experimental constraints for network inference. Here, RPPA-based phospho-proteomics data from the TCGA project for breast and ovarian cancers is used (43, 39). The utility and reliability of TCGA RPPA data as input for probabilistic graphical models to infer signaling networks have been previously reported and validated (36, 68). This previous work showed the ability of RPPA datasets to capture biologically relevant associations within individual cancer types. Each RPPA dataset includes 224 phospho and total proteins interrogated across 901 and 408 breast and ovarian cancer patients, respectively (see Table S1 for protein names).

##### Reference Network inference with the glasso algorithm

We have previously published a review and benchmark comparison of 13 established network inference methods in inferring protein-protein interaction networks from RPPA data (36). Therein, we benchmarked in their ability to reproduce interactions from the Pathway Commons database of molecular interactions as well as computational efficiencies. The benchmarking study also reported the precision and recall performances of an array of inference methods. Of our previously benchmarked regularized methods, graphical LASSO (glasso) is a method that balances computation speed (allowing scale up to larger systems) and performance measures (i.e., accuracy); additionally, glasso provides us a relatively simple framework by which prior information about molecular interactions from literature can be incorporated.

Using proteomic data, the Gaussian graphical model algorithm, glasso generates a partial correlation-based network (37). For this purpose, the algorithm estimates a sparse inverse of the covariance matrix, termed the precision matrix (*Θ*= *Σ*^−1^), using an L1 penalty. The precision (i.e., inverse covariance) matrix is an estimate through maximization of the log-likelihood function by applying the L1 penalty;

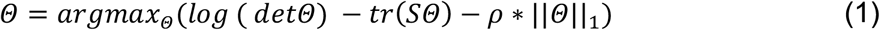

where S is the empirical covariance matrix and *ρ* is the weight of the complexity term.

Next, the precision matrix is converted to an undirected partial correlation network in which each interaction represents the direct associations between node pairs with no confounding contributions from other nodes. The edge strength (*W*_*ij*_) is the degree of partial correlation between node i and node j, and is estimated from the precision matrix using the equation

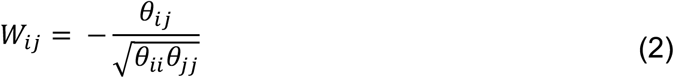

where *θ*_*ij*_ are elements of the estimated precision matrix 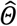 and *θ*_*ii*_ represents the auto-correlations (diagonal elements of the precision matrix). The resulting edge set with real-valued association (edge) strengths defines the reference network. The network models for breast and ovarian cancers are available in Tables S4 and S5 in a simple interaction format (SIF).

##### Prior-glasso algorithm

We have developed a modified version of the glasso algorithm, termed prior-glasso. The prior-glasso integrates experimental constraints with prior information from signaling databases as represented in the signedPC data source. This prior information is incorporated into the inference with a probabilistic term. The scheme favors prior interactions that conform to the experimental data, eliminates priors that do not conform with data, and predicts additional interactions that are not in databases but dictated solely by the experimental data. In order to enable comprehensive adaptive response quantification in subsequent steps, the proteins that are not covered by the patient RPPA (used for network building) but measured within the drug response data (used for TargetScore calculation) are also included solely based on prior information. The prior information narrows the potential solutions to a biologically more relevant domain of the search space by favoring the models that are enriched with known biology among solutions with similar errors. The signaling prior is represented with the information matrix *λ* which is a symmetric adjacency matrix of interactions with the entries

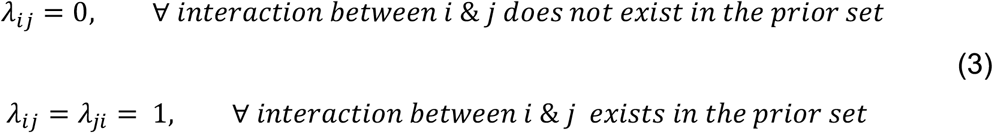

The prior information is integrated into the Gaussian graphical model as an additional term that modifies the log-likelihood equation in the form of (*κ*λ*_*P*P*_)∥*Θ*∥_1_ to the Equation 1. The precision matrix *Θ* is then estimated through the maximization of the modified log-likelihood function as in the glasso algorithm;

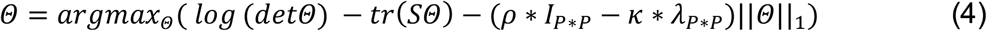

where *I* is the identity matrix with dimension *P*P, ρ* is the weight of the penalty parameter for complexity and *κ* serves as a scalar parameter for the contribution of the prior information matrix, P is the number of variables in the system. Here the parameters *ρ* and *κ* are tuned through a two-dimensional Bayesian information criterion (BIC) grid search. In BIC, the paired values of *ρ* and *κ* are explored to search for a parameter set with the smallest (i.e., more preferable) BIC (72)

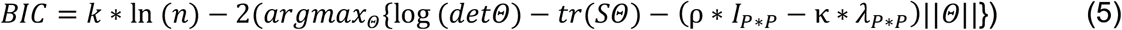

where *n* stands for the sample size of the data, *k* stands for the number of non-zero elements in the estimated precision matrix. BIC is calculated for every parameter combination through varying ρ and κ, ranging from 0 to 1 with step sizes of 0.02 in log10 space {10^−2^, 10^−1.98^, …, 10^−0.02^, 10^0^}. Optimized parameters *(ρ, κ)* were chosen at the lowest BIC value.

##### Edge directionality

Directions of the inferred edges are extracted from the signedPC pathway source. The edge direction is inferred into the reference network with an indicator function, 1_*B*_(*λ*_*ij*_)such that (1_*B*_(*λ*_*ij*_)= 0*or*1) based on the prior network in SignedPC. The below formulation leads to a partially directed network.

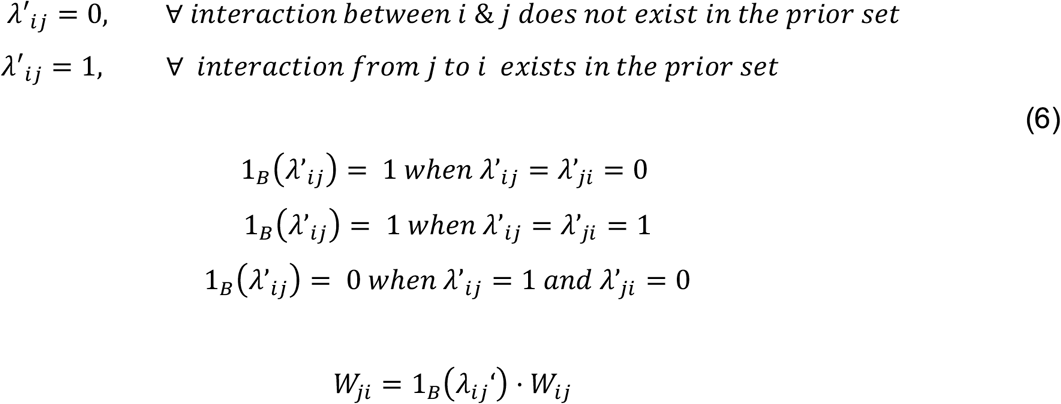

The *λ’*_*ij*_ represents the directional prior information from the signedPC resource (44). Equation 6 introduces the directions based on the prior information network and those interactions that are not covered in signedPC (data-driven interactions) remain bidirectional.

##### Network model robustness

To evaluate the robustness of the data-driven networks, we devised a bootstrapping scheme that involves the comparison of models inferred with partial vs. complete datasets. The models from partial data are generated using 1000 bootstraps (randomly sampling the partial data) with 50%, 60%, 70%, 80%, and 90% of the data. The similarity between the models from complete vs. partial data quantifies the robustness score. To measure the similarity, the partial correlation network is transformed into a binary network where edges have the values {*e*}_*ij*_ = {−1,0,1} such that -1 and +1 indicate presence whereas 0 indicates lack of interaction. The *e*_*ij*_ is the binary value of the edges between nodes *i* and *j* and it is assigned with a step function.

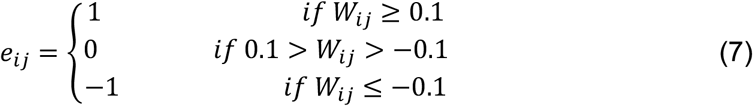

The sign represents the up (positive) vs. down (negative) regulation of the downstream node by the upstream. The weak edges with values that are close to 0 (|*W*_*i*j_|<0.1) are deleted from the model. Finally, the degree of robustness is quantified with a score that captures the overlap between the edges predicted by the partial data and edges predicted by the complete data.

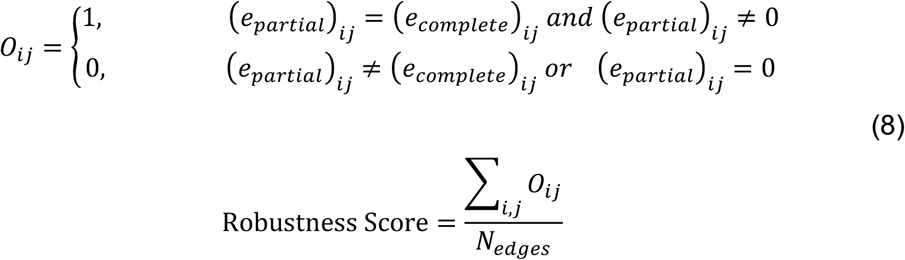

where, *O*_*ij*_ quantifies whether the non-zero edges between nodes *i* and *j* overlap with each in the models from complete vs. partial data. The *(e*_*partial*_)_*ij*_ is the binary value of the edge between nodes *i* and *j* in the model generated with the partial data. Similarly, *(e*_*complete*_)_*ij*_ is the edge value in the model generated with the complete data. *N*_*edges*_ is the number of edges in the model from the complete data. The sum of *O*_*ij*_ over all edges quantifies the total overlap. The ratio of the overlap between models to the total number of edges quantifies network robustness.

##### Reference network inference accuracy

We evaluate the model accuracy for both glasso and prior-glasso with 5-fold cross-validation. The cross-validation (CV) error is calculated based on the BIC with the log-likelihood evaluated. In the BIC, the precision matrix originates from the training set (80% coverage) and the empirical covariance from the validation set (20% coverage).

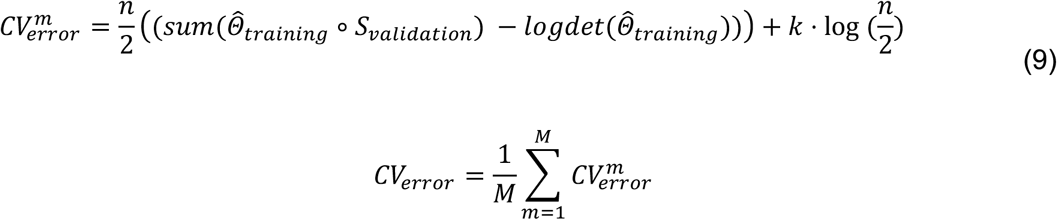

*Θ*_*training*_ is the estimated precision matrix from the training model, *S*_*validation*_ is the empirical covariance matrix generated from the validation data, *n* is the number of observations, *k* is the non-zero parameters estimated, *m* is the bootstrapping index. The training and validation set populations are generated with 500 bootstraps (*M*=500) from complete data without polluting the two sets with each other in individual bootstrapping steps. To quantify the model accuracy, the error is calculated for each bootstrapped training and validation data.

#### TargetScore calculation

TargetScore quantifies the collective adaptive pathway responses to a perturbation for each proteomic entity on the reference network. The TargetScore calculation requires at least a phospho-proteomic profile from a single sample under one perturbation condition as well as the unperturbed control condition. TargetScore is calculated as the sum of the response from each phospho-protein level and its pathway neighborhood. The calculation combines the cell type-specific drug response data with the pathway neighborhood information as encoded in the reference network model. The mathematical formulation of TargetScore is

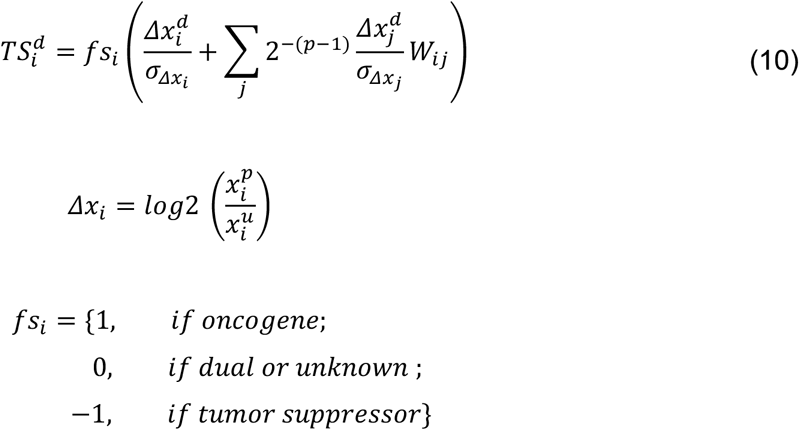

where *fs*_*i*_ represents the function score, *Δx*_*i*_ is the proteomic response, which is log normalized with respect to the matching readout from the unperturbed conditions. *σ*_*i*_ is the standard deviation of *Δx*_i_ over all samples for each protein entity across all conditions (e.g., culturing conditions, time points, drug doses). The optional step involving normalization of readouts with respect to the standard deviation enables the algorithm to scale the readouts to a comparable dynamic range for each antibody and cross-sample comparisons. Node *j* is a node in the pathway neighborhood of node *i*, with readout *Δx*_*j*_ and standard deviation, *σ*_*j*_. *p* is the pathway distance between the nodes *i* and *j*. The term 2^(p-1)^ ensures the dissipation of pathway influences from high order pathway neighborhoods. Note that, *P*_*max*_ is set to 1 (limiting the calculations to the first neighborhood of interactions) in the current implementation but it can be adjusted as needed. *W*_*ij*_ represents the signaling interaction between nodes *i* and *j* as inferred within the reference network. The extremely weak partial correlations corresponding to {-0.05 < W_ij_ < 0.05} are removed to reduce noise and {W_ij_} are normalized with respect to the maximum absolute value of W_ij_ to ensure resulting {W_ij_} is in [-1,1] boundary. The removal of the edges with very weak W_ij_ enables TargetScore calculations with a more interpretable reference network and minimal impact on the outcome. The cumulative TargetScore over all doses (ds) is formulated as

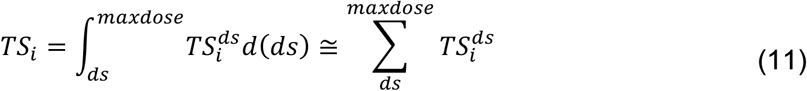

*TS*_i_ gives the cumulative TargetScore across all doses yet a dose-dependent TargetScore can also be analyzed.

##### Function Score

The optional parameter, function score, which annotates the oncogenic and tumor suppressor roles, is assigned to each of the proteomic entities measured within the RPPA platform. The oncogenic role for each measured entity is extracted from the COSMIC (version 89) (69). The resulting function scores were refined with literature-based, manual curation. A functional score of +1 is assigned to proteomic entities representing total level and activating phosphorylation of oncogenes or deactivating phosphorylation of tumor suppressors. Similarly, a functional score of (−1) is assigned to total levels and activating phosphorylation of tumor suppressors and inhibitory phosphorylation of oncogenic proteins. The activating vs inhibitory roles of phosphorylation are assigned based on information on the Phosphosite database or manual curation (70). The function scores are available at https://github.com/korkutlab/targetscore as part of the TargetScore package and in Table S3.

##### Statistical assessment of TargetScore

A potential caveat in the TargetScore calculations is the bias introduced by the hyper-connected hub nodes. Such hyper-connectivity can lead to artificially high TargetScores solely driven by the network topology and independent of the context-specific drug response data. To eliminate the connectivity bias, we assess the significance of TargetScore for each proteomic entity. For this purpose, the probability of observing a TargetScore is calculated over a fixed reference network structure and randomized drug response data (randomized protein labels). The randomized data is generated by sampling from the proteomic responses across all conditions for each antibody and assigning random labels to sampled data. Based on calculations with 1000 randomly bootstrapped data sets, the null distribution of TargetScores for a given network topology is calculated. Next, the FDR-adjusted q-value is calculated for the *TS*_*i*_ for each node (*i*) based on the Benjamini-Hochberg method. Adaptive responses with high TargetScore values and low q-value are nominated for further analysis.

##### TargetScore module rank and candidate selection

To identify adaptive response modules and mechanisms of action modules, we map the top 10% of proteins with the highest TargetScore values to the underlying reference networks and detect the highly scored multi-molecule modules. Subnetworks with high TargetScore values are identified as adaptive resistant responses modules while the subnetworks with the lowest TargetScores are nominated as potentially associated with mechanisms of action. Potential adaptive resistance pathways (or modules) are detected through differential analysis of TargetScores for each protein in drug-resistant versus sensitive samples. The comparison can be made by one-to-one analysis of sensitive and resistant samples or with a t-test between resistant and sensitive populations when a sufficient number of samples exists.

## Experimental Methods

### Reagents, cell lines, cell cultures

MCL1 inhibitor-S63845 was purchased from MedChemExpress. BCL2 inhibitor (ABT199), SHP2 inhibitor (SHP099) and PARP inhibitor (AZD2281) were purchased from Sellekchem. The compounds were prepared as stock solutions in DMSO. The drugs were stored at -20°C and diluted immediately before use. HCC1954 and SKOV3 cell lines were from the MD Anderson Cancer Center cell line repository and had no more than 10 passages. The cells were thawed and cultured in RPMI 1640 supplemented with 10% FBS two weeks before experiments. The cells were cultured at 37°C in a humidified atmosphere with 5% CO2.

### Cell proliferation assays

Cell proliferation was measured according to the instructions of the PrestoBlue Cell Viability Assay kit (A13261, Life Technologies). 2×10^3^ cells were seeded triplicate into 96-well plates for culturing overnight. The next day, cells were treated with either vehicle or kinase inhibitors.72 hours later, cells were collected and measured with a SYNERGY H1 microplate reader and analyzed using the Gen5 software (BioTek).

### Induction of MCL1 and BCL2 inhibitor-resistance in HCC1954 cells

HCC1954 cells were thawed and passaged two times in a medium of RPMI 1640 supplemented with 10% FBS. The cells were seeded into 10 cm culture plates and changed fresh medium plus MCL1 and BCL2 inhibitors of increasing concentrations (from 0.1, 0.2, 0.4, 0.6, 0.8, 1.0, 1.5, 2.0, 2.5, 3.0, 3.5, 4.0 µm of inhibitors). Incubating for ∼4 months under inhibitor stress, cells were diluted and seeded in new 10 cm culture plates. Cells were continued to culture in the medium plus MCL1 and BCL2 inhibitors for colony formation in the long term. The single colonies were picked and cultured in fresh medium. Both the mixture and the single colonies were collected and frozen down at liquid nitrogen.

### RPPA (Reverse Phase Protein Array)

The cells were washed 3 times with cold PBS and then suspended in an RPPA buffer supplemented with proteinase and phosphatase inhibitors (Pierce, Rockford, IL, USA). The cell suspension was vortexed for 15 seconds, placed on an end-over-end rotator for 30 min at 4°C and centrifuged at 14,000 x g for 15 min at 4°C. The lysates were prepared to provide 1-1.5mg/ml of total protein lysate. RPPA analysis samples were prepared by adding SDS Sample Buffer, α-mercaptoethanol and RPPA Working Solution to obtain a final concentration of 0.5mg/ml. Samples were heated for 8 min at 100°C, centrifuged 2 min at 14,000 x g and stored at -80°C. The RPPA was performed at the MD Anderson Cancer Center Functional Proteomics core facility.

### Growth rate and drug synergy calculations

GR estimates the growth rate inhibition using endpoint or time-course assays by eliminating the confounding impact of doubling times in cell viability measurements (40). The GR value is the ratio between growth rates for the treated and untreated conditions normalized to number of cell divisions. A GR value greater than 1 indicates increased growth, values between 0 to 1 indicates cytostatic effects whereas GR < 0 indicates cytotoxicity. The GR metric is quantified as:

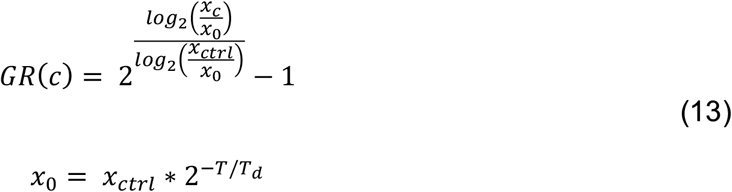

*T*: The duration of the assay, *c*: drug concentration, *T*_*d*_: doubling time, *x*_*c*_: cell count at time t, *x*_*ctrl*:_ control sample cell count, *x*_*0*:_ the cell counts from a sample with no drug treatment grown in parallel and measured just before drug exposure. The *x*_*0*_ can be estimated from the previously measured (or literature extracted) *T*_*d*_.

